# Enhancers that regulate *TNF* gene transcription in human macrophages in response to TLR3 stimulation

**DOI:** 10.1101/2022.06.13.496019

**Authors:** Junfeng Gao, Yapeng Li, Dianzheng Zhao, Xiaoyu Guan, Kirby Motsinger, James Scott-Browne, Hong Wei Chu, Hua Huang

## Abstract

Macrophages play a critical role in inflammatory responses during infections. These cells are activated by infections through stimulation of TLRs expressed on their cell surface and produce pro-inflammatory cytokines, including TNF. However, distal enhancers that regulate *TNF* gene transcription in human macrophages have not been investigated. This study used an unbiased genomic approach to identify six candidate enhancers in human primary alveolar macrophages within a 131 kb region from the transcription start site (TSS) of the *TNF* gene, covering 13 genes. Of these candidate enhancers, five showed enhancer activity, with three targeting the *TNF* gene and two targeting neighboring genes. Deletion of the distal *TNF* E-16 enhancer led to a 73% reduction in *TNF* gene transcription in response to poly (I:C) stimulation in the THP-1 human leukemia monocytic cell line. Additionally, deletion of the E-7.1/hHS-8 enhancer resulted in a 41% reduction in *TNF* mRNA, while deletion of the PE enhancer had a lesser effect, resulting in a 52% reduction in *TNF* gene transcription. Massively parallel reporter assays (MPRA) indicated that the transcription factor AP-1 and EGR1-binding sites at the distal *TNF* E-16 enhancer were crucial in mediating enhancer activity. This study shows that both distal and proximal enhancers work together to fully transcribe the *TNF* gene in human macrophages in response to TLR ligand poly (I:C) stimulation.

## Introduction

Macrophages are among the first immune responders to pathogens (1,2), and play a pivotal role in inflammatory responses. They respond to infections through pattern-recognition receptors, including Toll-like receptors (TLRs) (3,4). TLRs bind to viral and bacterial productions derived from many bacteria and viruses at some point of their replication cycle (5,6). TLR3 can induce immune response against double-stranded RNAs produced by viral infection. Several synthetic dsRNAs are reported to activate TLR3, including polyriboinosinic: polyribocytidylic acid (poly(I:C)) (5,7,8). TLR activation induces the production of chemokines and cytokines in macrophages (9,10). However, overproduction of these cytokines and chemokines can lead to tissue damage during severe infections (11,12) and severe toxic effects in cancer immune therapies (13).

Despite the critical role of TNF-α in the immune response, the transcriptional codes that regulate *TNF* gene expression in response to TLR stimulation and infected epithelial cells are still poorly understood. Enhancers, which are segments of DNA located in the non-coding regions of genes (14), play a critical role in regulating gene expression. In this study, we focus on enhancer regulation of the *TNF* gene in human lung macrophages.

Locating critical enhancers can be a significant challenge because they can be located up to 100kb from the transcription start sites in non-coding regions that make up 99% of a genome (15). The *TNF* gene locus lies in mouse chromosome 17 and human chromosome 6 and is comprised of the *TNF* gene and the genes encoding lymphotoxin-a and lymphotoxin-b (LTA and LTB) (16). Some enhancers have been reported to regulate *TNF* gene transcription in mouse and human macrophages. A distal enhancer element 9 kb upstream of the mouse *Tnf* mRNA cap site (HHS-9) can bind transcription factor NFATp and participate in intrachromosomal interactions with the *Tnf* promoter in mouse T cells upon activation (17), while a distal hypersensitive site ∼8 kb upstream of the human *TNF* transcription start site (hHS-8) is required for and mediates IFN-γ-stimulated augmentation of LPS-induced *TNF* gene expression in macrophages via binding of IRF1 to a cognate hHS-8 site in human monocytes/macrophages (18). Additionally, hHS-8 coordinately regulates the *TNF* and *LTA* gene expression in activated human T cells via a discrete and highly conserved NFAT binding site (19). However, how the *TNF* gene is regulated by enhancers in human macrophages is not completely understood. One challenge is to find target genes for enhancers. Enhancers often work from a distance and cross genes. The bioinformatics approach often assigns enhancers to the nearest genes (15). This approach often misses distal enhancers. Although the chromatin capture assay is informative in finding distal enhancers (20), this assay does not provide information regarding the relative contribution of enhancers to target gene transcription.

We identified six candidate *TNF* enhancers within 131 kb of the TSS of the *TNF* gene using H3K27ac ChIP-seq and ATAC-seq in primary human alveolar macrophages. Proximal enhancer (PE), E-33, E-16, E-7.1, and E+85 exhibited enhancer activity in reporter gene assays. Deletion of the distal *TNF* E-16 enhancer resulted in a 73% reduction in *TNF* gene transcription in the matured human leukemia monocytic cell line THP-1 in response to poly (I:C) stimulation. Deletion of the PE enhancer resulted in a 52% reduction in *TNF* gene transcription to a lesser extent. Mutational analysis using massively parallel reporter assays (MPRA) revealed that transcription factor AP-1 and EGR1-binding sites at the distal *TNF* E-16 enhancer were critical in mediating the E-16 enhancer activity. Our study demonstrates that both distal and proximal enhancers work together to fully transcribe the *TNF* gene in human macrophages upon TLR ligand poly (I:C) stimulation.

## Results

### Identification of candidate *TNF* enhancers that respond to poly (I:C) stimulation

To determine the time course for *TNF* mRNA expression in human alveolar macrophages (AMs) in response to poly (I:C) stimulation, we treated primary human AMs from 7 donors without or with poly (I:C) for 4, 8, or 24 hours. We observed that *TNF* mRNA started to increase 4 hours after stimulation and reached the highest levels at 8 hours after stimulation (104.8-fold compared with the *TNF* mRNA in resting human primary AMs) and remained 15.6-fold higher than that in resting cells at 24 hours after stimulation (Fig. 1).

**Figure 1.**
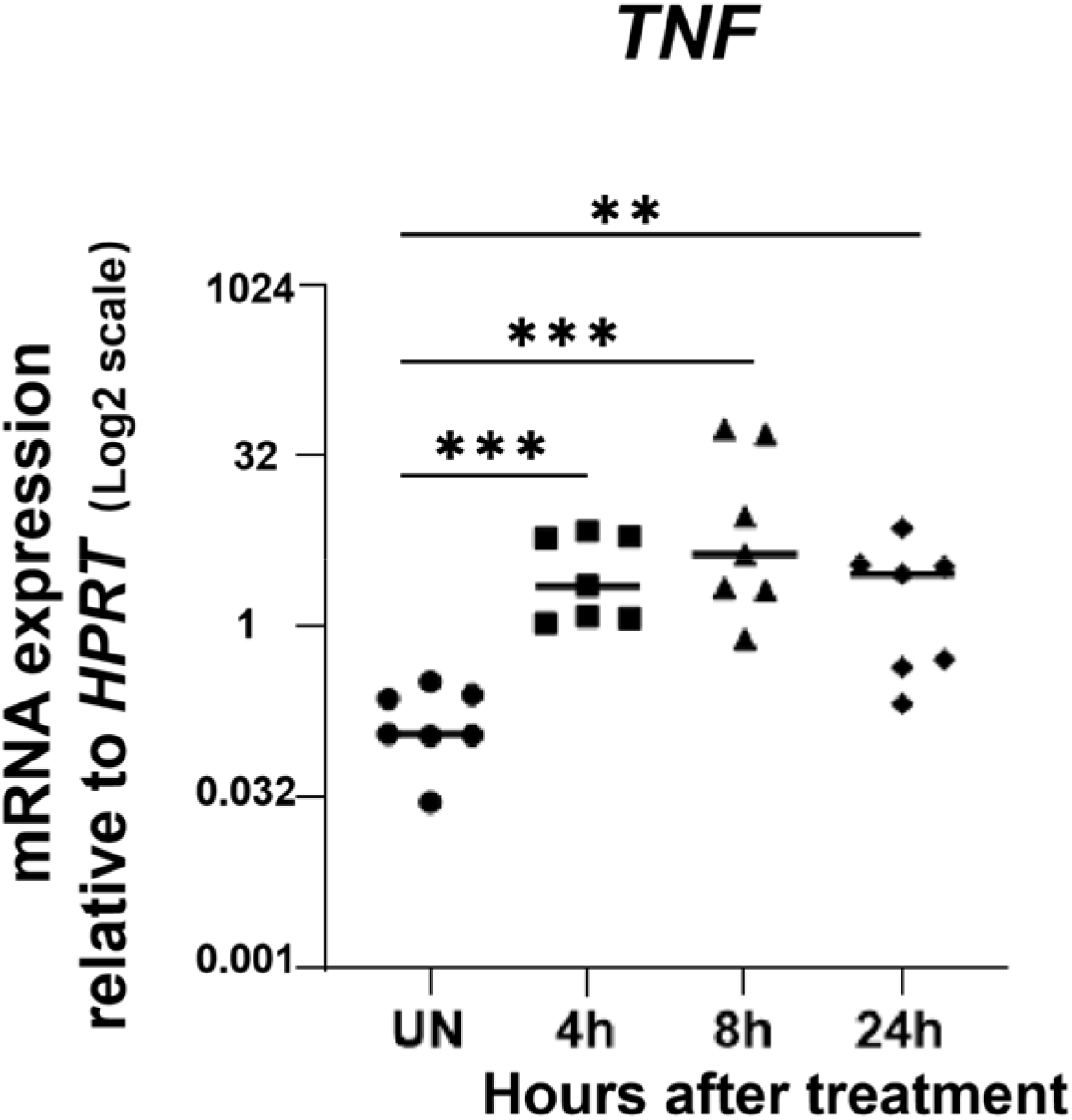
Primary human alveolar macrophages transcribe *TNF* genes to extremely high levels in responding to poly (I:C) stimulation. Human alveolar macrophages were untreated (UN), treated by poly (I:C) for four hours (4h), eight hours (8h) or twenty-four hours (24h). qPCR analysis of the mRNA expression of *TNF* genes. *P* values were calculated using the Mann-Whitney U test. Data represent mean ± SEM of seven biological samples. ** *P* < 0.01; *** *P* < 0.001.

Based on the time-course experiment, we chose 4 hours of treatment to detect early changes in H3K27ac histone modification and chromatin accessibility. We treated AMs without or with poly (I:C) for four hours. We performed H3K27ac ChIP-seq and Omni-ATAC-seq to identify non-coding DNA regions associated with increased H3K27ac modification and chromatin accessibility. We showed that there were six potential enhancer regions that were associated with increased chromatin accessibility and H3K27ac modification within 131 kb of the human *TNF* gene (−42.6 upstream and +88.8 kb downstream from the TSS of the *TNF* gene), which covers the intergenic regions between 13 genes, including *LTA, LTB, DDX39B, GPANK1*, etc. (Fig. 2). We excluded *BAG6* exon (Fig. 2). We named these candidate enhancers proximal enhancer (PE, minus the core promoter), E-33, E-16, E-7.1, E+44 and E+85 based on the distances of the candidate enhancers to the transcription start site (TSS) of the *TNF* gene.

**Figure 2.**
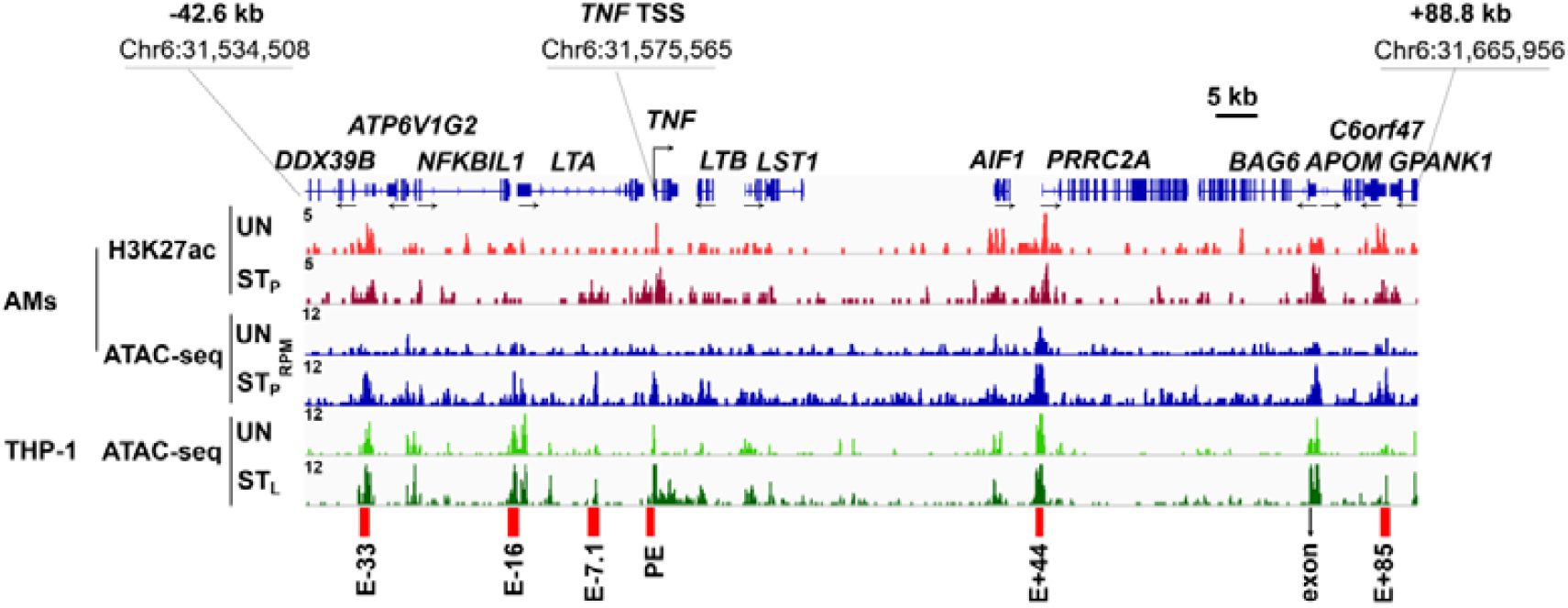
Identification of the candidate *TNF* enhancers. Representative Integrated Genome Viewer (IGV) tracks from H3K27ac ChIP-seq and Omni-ATAC-seq. Human alveolar macrophages (AMs) with unstimulation (UN) or poly (I:C) stimulation (ST_P_) for four hours were used for H3K27ac ChIP-seq and Omni-ATAC-seq. The ATAC-seq data from THP-1 cells, which were either unstimulated (UN) or stimulated with LPS (ST_L_), were obtained from the Gene Expression Omnibus (GEO) database with accession number “GSE147314”. Red bars indicate candidate *TNF* enhancers that show increased H3K27ac modifications and chromatin accessibility. RPM: reads per million mapped reads; E: enhancer; PE: proximal enhancer. The numbers following E indicate the distance (kb) of enhancer to the TSS of the *TNF* gene; + means the enhancers are located at the downstream of the TSS, and – means the enhancers are located at the upstream of the TSS. One IGV track was from one biological sample, representing two biological replicates with similar patterns.

### The *TNF* E-16, PE, E-33, E-16, E-7.1, and E+85 enhancers possess enhancer activity

Not all non-coding regions associated with H3K27ac modification and increased chromatin accessibility possess enhancer activity. Primary human AMs are difficult to transfect with electroporation or transduce with lentivirus, we chose the human monocytic cell line THP-1 cells as a cellular model to investigate the role of various candidate enhancers in transcribing the *TNF* gene in response to poly (I:C) stimulation. THP-1 cells can be matured with phorbol 12-myristate 13-acetate (PMA) to macrophages to transcribe cytokine genes at the levels comparable to macrophage (21,22). The matured THP-1 cells transcribed 11-fold more *TNF* mRNA than immature THP-1 cells in response to poly (I:C) stimulation (Fig. S1). We further verified chromatin accessibility in LPS-stimulated matured THP-1 cells were strikingly similar to those of poly (I:C)-stimulated AMs (Fig. 2), indicating that matured THP-1 cells were suitable for functional analysis of the candidate enhancers. ATAC-seq data of THP-1 cells were downloaded from the Gene Expression Omnibus (GEO): “GSE147314”) (23). To access the enhancer activity of the potential enhancers, we refined the published MPRA (24) to allow the cloning of longer PCR-amplified enhancer fragments and sequencing barcode DNA and RNA using the Nova-seq multiplexing platform. We cloned the PCR products of the candidate enhancers (366-582 bp) into the LentiMPRA vector containing a minimal promoter and manually selected barcoded *eGFP* reporter gene (24) (Fig. 3A). We used a non-coding DNA fragment that was not associated with H3K27ac or chromatin accessibility as non-enhancer (NE) control. We transduced THP-1 cells with the recombinant lentivirus containing the candidate enhancers. We matured the transduced THP-1 cells with PMA for three days. RNA and DNA barcodes in RNA and DNA samples prepared from transduced cells were sequenced to determine the number of RNA barcode transcripts and DNA inserts. The ratios of barcode RNA transcripts to barcode DNA inserts (as input controls) were used to determine enhancer activity. We found that the *TNF* E-33, E-16, E-7.1, PE and E+85 showed significant enhancer activity (Fig. 3B).

**Figure 3.**
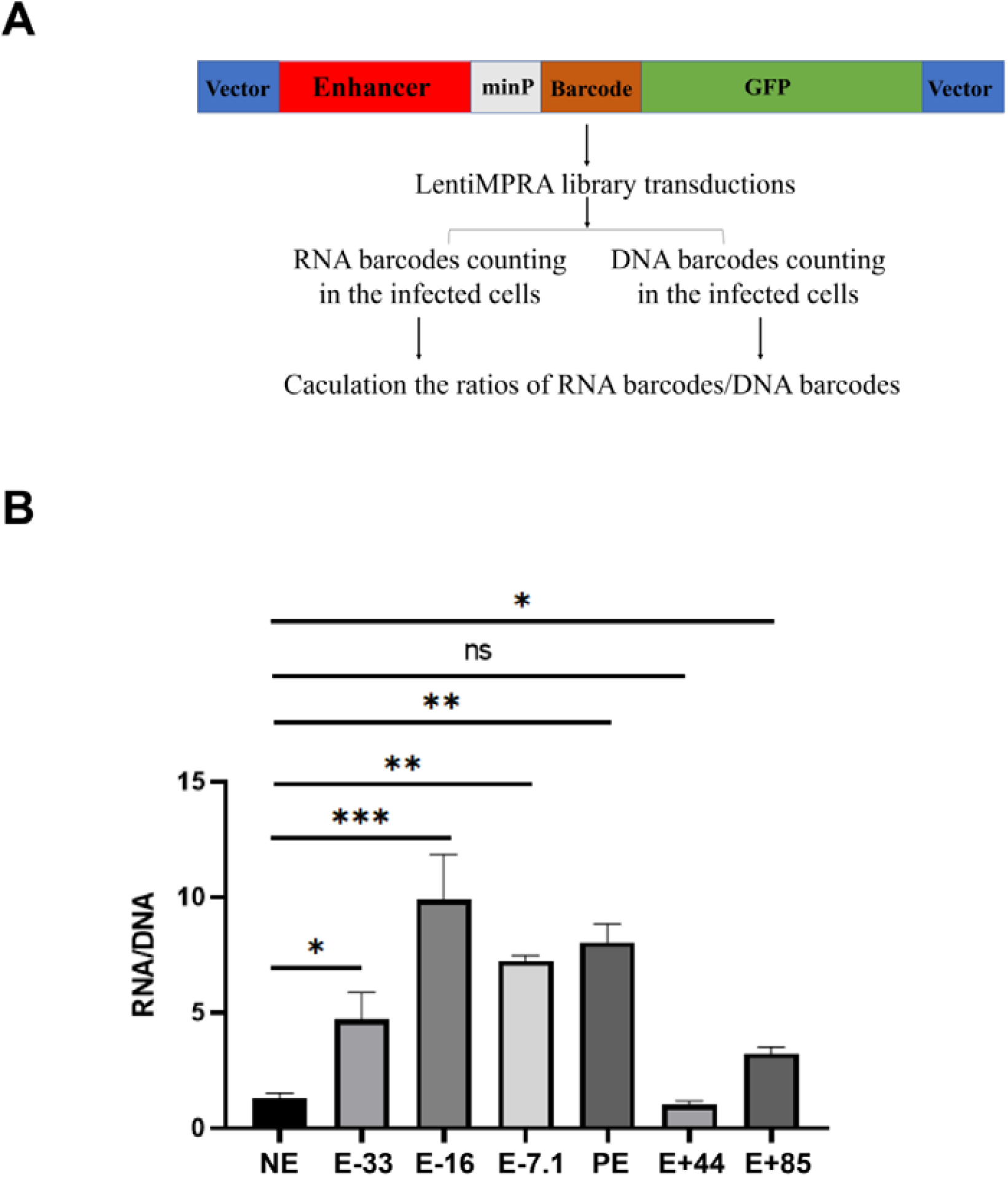
*TNF* E-33, E-16, E-7.1, PE and E+85 possess enhancer activity in response to poly (I:C) stimulation. **A**. Barcoded GFP reporter gene constructs. **B**. qPCR analysis of RNA barcodes or DNA barcodes in samples prepared from the enhancers-transduced-THP-1 cells. Data represent mean ± SEM of three transduced samples from three independent experiments. *P* values were calculated using two-tailed student’s *t* test. * *P* < 0.05; ** *P* < 0.01; *** *P* < 0.001.

### The *TNF* E-16, E-7.1 and PE enhancers significantly contribute to *TNF* gene transcription in the context of chromatin in response to poly (I:C) stimulation

The reporter gene assay does not measure gene transcription in the context of naïve chromatin and does not provide information on target gene. To determine whether the candidate enhancers that possessed enhancer activity target the *TNF* gene and the contribution of these enhancers to *TNF* gene transcription in response to poly (I:C) stimulation, we deleted the PE, E-33, E-16, E-7.1 and E+85 enhancers using our improved CRISPR/Cas9 deletion method. We chose NE as negative control. The current CRISPR deletion method using two sgRNA guides often resulted in deletions occurring in one chromosome, creating heterozygous deletion that does not have a phenotype. To overcome this technical challenge, we targeted each enhancer with four sgRNA guides, each contained within a bicistronic gene co-encoding for a different fluorescence protein GFP, RFP, BFP, or Thy1.1 molecule (Fig. 4A). We found that 9.4 % of THP-1 cells transduced with lentivirus containing the four sgRNA guides expressing BFP, GFP, RFP, and Thy1.1 (Fig. 4B). FACS-sorted cells positive for BFP, GFP, RFP, and Thy1.1 achieved complete homozygous deletion of NE, E-33, E-16, E-7.1, PE and E+85 enhancers in bulk using this newly improved method (Fig. 4C). Deletion of the E-16 enhancer resulted in a 73 % reduction in *TNF* mRNA expression, deletion of the E-7.1 enhancer led to a 41 % reduction in *TNF* mRNA expression, deletion of the PE enhancer led to a 52 % reduction in *TNF* mRNA expression, deletion of the E-33 and E+85 did not affect *TNF* mRNA expression although deletion of the E-33 reduced *DDX39B* by 41 % and deletion of E+85 reduced *C6orf47* by 66 %, reduced *GPANK1* by 47 % (Fig. S2). In contrast, deletion of the E-16 or the PE did not affect *LTA* and *LTB* mRNA expression (Fig. 4 E and F). These data demonstrate that E-16 is critical in *TNF* gene transcription in response to poly (I:C) stimulation.

**Figure 4.**
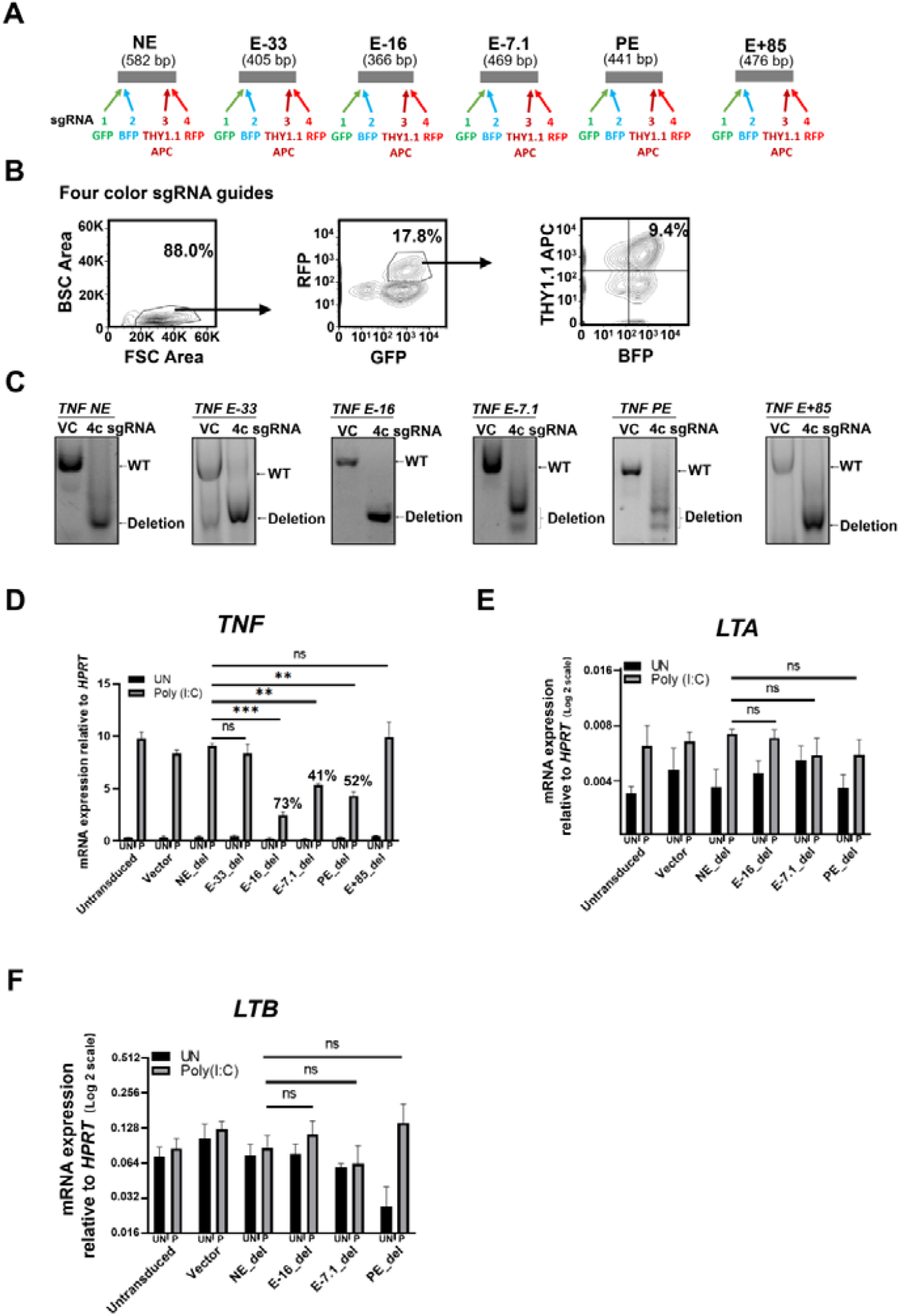
The *TNF* E-16 is required for *TNF* gene transcription in response to poly (I:C) stimulation. **A**. Targeting one enhancer with bicistronic sgRNA guides co-expressing BFP, GFP, RFP, and Thy1.1. **B**. FACS sorting gates. THP-1 cells were transduced with BFP, GFP, RFP, and Thy1.1 sgRNA guides (4c sgRNA guides). BFP^+^, GFP^+^, RFP^+^, and Thy1.1APC^+^ transduced cells were FACS-sorted using sorting gates indicated. **C**. DNA deletion efficiency analysis. Vector control (VC) and deleted DNA fragments in bulk FACS-sorted BFP^+^, GFP^+^, RFP^+^, and Thy1.1APC^+^ cells were analyzed with PCR. **D**. *TNF* mRNA expression in the FACS-sorted transduced cells was measured by qPCR. The percentages indicate the percent reduction in *TNF* mRNA expression in enhancer-deleted (del) relative to non-transduced or vector-transduced THP-1 cells. **E**. *LTA* mRNA expression in the FACS-sorted cells was measured by qPCR. **F**. *LTB* mRNA expression in the FACS-sorted cells was measured by qPCR. P: poly (I:C). *P* values were calculated using two-tailed student’s *t* test. Data represent mean ± SEM of four transduced samples from two independent experiments. * *P* < 0.05; ** *P* < 0.01; *** *P* < 0.001.

### The AP-1 and EGR1 binding sites are essential for driving the E-16 enhancer activity

To locate the poly (I:C) responsive TF binding sites within the E-16 enhancer, we used Massively parallel reporter assays (MPRA) with randomly selected barcodes to measure enhancer activity of E-16. We found that the E-16_1 but not E-16_2 showed significant enhancer activity (Fig. 5A). To determine which TF binding sites in the E-16_1 fragment drive enhancer activity, we performed scan mutation analysis. In the scan mutants we replaced WT sequences 20 bp at a time with sequences (GTCAAGAGGTCGAGTCAAGA) that did not contribute to transcriptional activity (25). We found that scan mutant 3 and scan mutant 6 significantly lost their ability to enhance barcoded *GFP* reporter gene transcription (Fig. 5B). We analyzed TF binding sites in the scan mutant 3 and scan mutant 6 and found that EGR1 binding site in the scan mutant 3 and AP-1 binding site in the scan mutant 6 (Fig. 5C). Thus, these results demonstrate that EGR1 and AP-1 binding sites at the *TNF* E-16 enhancer are required for the E-16 enhancer activity.

**Figure 5.**
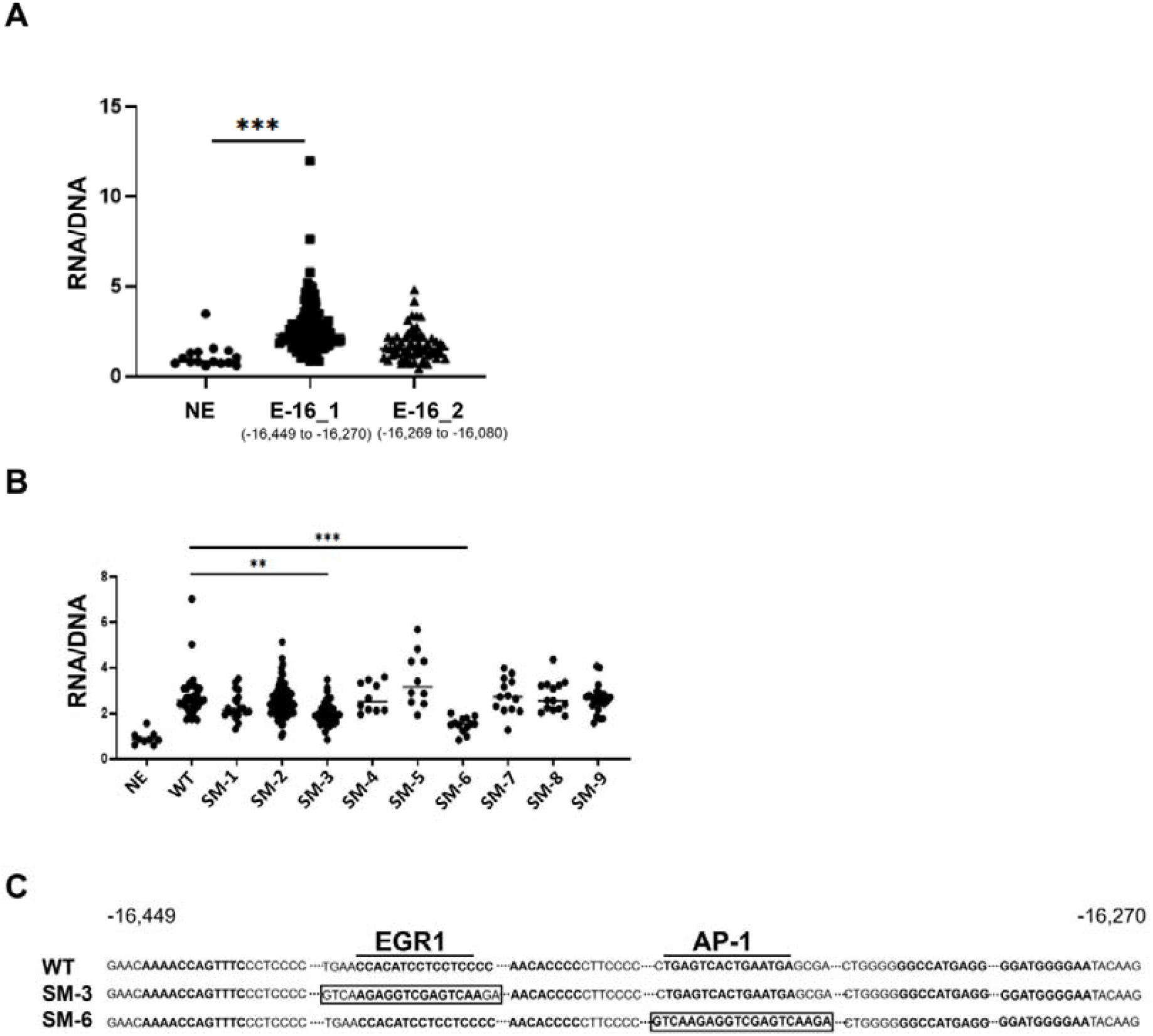
The AP-1 and EGR1 binding sites are essential for driving the E-16 enhancer activity. **A**. Enhancer activity analysis of the fragments from position - 16,449 to -16,270 and -16,269 to -16,080 of the E-16 enhancer. The ratios of barcode RNA transcripts to barcode DNA inserts (as input controls) were used to determine enhancer activity. Each point represents one barcoded GFP mRNA expression normalized against the barcoded GFP DNA insertion pooled from three transductions from three experiments. **B**. Scan mutational analysis of the fragment from position - 16,449 to -16,270 of the E-16 enhancer. **C**. Mutated EGR1 and AP-1 binding sites are shown. Mean ± SEM were calculated from three transductions from three experiments. *P* values were calculated using the Mann-Whitney U test. ** *P* < 0.01; *** *P* < 0.001.

## Discussion

Our study addresses the long-standing problem of assigning enhancers to target genes. We improved the CRISPR deletion method and used it to delete five candidate enhancers we identified in the 131 kb region consisting of 13 genes, including the *TNF* gene. Of these candidate enhancers, three targeting the *TNF* gene and two targeting neighboring genes. We demonstrate that both distal and proximal *TN*F enhancers work together to fully transcribe the *TNF* gene in human macrophages upon TLR ligand poly (I:C) stimulation.

Given the difficulty of transfecting primary alveolar macrophages, we opted to use THP-1 monocytic cell line in our functional analysis of candidate enhancers. Our results showed that THP-1 monocytes, upon maturation into macrophages, displayed a much higher capacity to transcribe cytokine genes in response to poly (I:C) stimulation, and had chromatin accessibility regions similar to those of alveolar macrophages following TLR stimulation. We also discovered that not all DNA regions associated with increased H3K27ac modification and chromatin accessibility had enhancer activity, and not all DNA regions with enhancer activity contributed to *TNF* gene transcription in the context of chromatin.

Of the enhancers we identified, PE, E-7.1, and E-16 were the most significant contributors to *TNF* gene transcription in response to poly (I:C) stimulation. While the E-16 enhancer is located between the *LTA* and *LTB* genes, deletion of this enhancer resulted in a profound 73% reduction in *TNF* gene transcription response to poly (I:C) stimulation, whereas deletion of this enhancer did not significantly alter the mRNA expression of *LTA* or *LTB* gene. The traditional mutational analysis is tedious. We refined MPRA to allow us to manually or randomly select barcodes to associate with enhancers of interest. We used manually selected barcodes to associate with longer PCR products that exceed the lengths that Ilumina instrument can sequence and used randomly selected barcodes to associate with 200-bp enhancers that contain WT and mutated TF binding sites. Our MPRA analysis further revealed that EGR1 and AP-1 binding sites were critical in the *TNF* E-16 enhancer activity.

We also identified E-7.1 within the previously reported hHS-8 enhancer with 458 bp overlap. The hHS-8 enhancer is required for IFNγ-primed macrophages to transcribe the *TNF* gene in response to LPS stimulation (18). This hHS-8 is also required for *TNF* and *LTA* gene expression in T cells (18,19). While hHS-8 was identified with traditional DNase Hypersensitivity (DHS) mapping, we identified E-7.1 with ATAC-seq, and its peak was narrower compared to the hHS-8 DHS peak. Deletion of the E-7.1/hHS-8 resulted in a 41% reduction in *TNF* mRNA, confirming its role in *TNF* gene transcription. We also determined the contribution of the proximal enhancer (minus core promoter) PE in *TNF* gene transcription in response to poly (I:C) stimulation. However, the E-33 and E+85 enhancers possessed enhancer activity but did not target the *TNF* gene. Instead, the E-33 enhancer targeted *DDX39B* gene located upstream of the *TNF* gene, whereas the E+85 enhancer targeted *C6orf47* and *GPANK1* located downstream of the *TNF* gene. We analyzed the published Hi-C data (GEO: GSE103477) (26,27) and found that high levels of interaction between *TNF* E-16, E-7.1, and PE in matured THP-1 cells under both resting and IFNβ-stimulated conditions. Although the chromatin capture assay is informative in determining enhancer and core promoter interactions (20), our investigation showed that in situ enhancer deletion is the most effective way to determine which enhancers target the *TNF* gene and the contribution of enhancers to *TNF* gene transcription.

Based on our findings, we propose that the E-16 works as an anchoring enhancer that interacts with the *TNF* core promoter, PE, and the E-7.1/hHS-8 enhancer. Disrupting the anchoring *TNF* E-16 would interfere with the formation of the *TNF* transcriptional complex. In contrast, disrupting other additive PE or E-7.1/hHS-8, which would not interfere with the interaction between the *TNF* core promoter and the anchoring *TNF* E-16 enhancer, causes a lesser reduction in *TNF* gene transcription in response to poly (I:C) stimulation. Our results demonstrate that that the PE, the E-7.1/hHS-8, and the E-16 contribute to full *TNF* gene transcription in response to poly (I:C) stimulation.

In summary, understanding the mechanisms of *TNF* gene regulation can provide mechanistic insights into the complex transcriptional regulation of the *TNF* gene in response to TLR3 stimulation. By identifying and characterizing the specific enhancers that contribute to *TNF* gene transcription, our study also provides potential molecular targets for medical interventions that target TNF-related diseases.

## Experimental procedures

### Human alveolar macrophages

Human lungs were obtained from de-identified male and female organ donors whose lungs were not suitable for transplant and were donated for medical research. We obtained the donor lungs through the International Institute for the Advancement of Medicine (Edison, NJ) and the National Disease Research Interchange (Philadelphia, PA). Research on these human lungs has been deemed as nonhuman subject research and is given IRB exemption because the donors are deceased and de-identified. Alveolar macrophages (AMs) were isolated from lavage of the lung before the instillation of elastase, as described previously (28). The purity of the AMs was 92.6 ± 2.8% as measured by immunostaining of cytocentrifuge preparations. AMs were frozen in a liquid nitrogen tank. Previous studies have compared freshly isolated and frozen AMs and did not find noticeable differences in their production of pro-inflammatory cytokines (28). AMs were cultured in DMEM (SH3024301, Thermo Fisher Scientific) plus 10% FBS, 100 units/mL penicillin, 100 μg/mL streptomycin, 2.5 μg/mL amphotericin B in the presence of 50 ng/ml GM-CSF (PeproTech, 300-03) in a humidified 37°C, 5% CO2 incubator. AM samples with high *TNF* mRNA expression without poly (I:C) stimulation were excluded from this study.

### Chromatin Immunoprecipitation and ChIP-seq

Chromatin immunoprecipitation was performed according to the published method (29). Briefly, human macrophages (5×10^6^) were not treated or treated with 1 μg/mL poly (I:C) for four hours were fixed with 1% formaldehyde (PI28908, Thermo Fisher Scientific), sonicated by using the Covaris S220 Focused-ultrasonicator in the SDS lysis buffer (1% SDS, 10 mM EDTA, 50mM Tris.HCl pH8) and precleared with Protein A Beads at 4 °C for 1h according to established protocols. The samples were incubated with 10 μg of following antibodies (1:100 dilution): anti-H3K27ac antibody (ab4729, Abcam, Abcam, Cambridge, MA) at 4°C overnight and then with protein A agarose/salmon sperm DNA slurry (Millipore, Cat# 16-157) at 4°C for 1h. The beads were washed and eluted as described. The crosslinking of eluted immunocomplexes was reversed and the recovered DNA was recovered using a QIAGEN QIAquick PCR purification kit (Qiagen, Valencia, CA). ChIP-seq library was prepared using TruSeq ChIP Library Preparation Kit (IP-202-1024, Illumina, San Diego, CA) according to the manufacturer’s instructions. Briefly, 10 ng of ChIPed DNA was converted into blunt-ended fragments. A single adenosine nucleotide was added to the 3’ ends of the blunt-ended fragments before ligation of indexing adapters to the adenylated 3’ ends. The ligated products were purified, size-selected and PCR amplified according to the manufacturer’s instructions. The quality and quantity of the DNA library were assessed on 4150 TapeStation System (Agilent, CA). Paired-ended sequencing was performed on an Illumina NovaSEQ6000 platform.

### Omni-ATAC-seq

Omni-ATAC-seq was performed according to the published method (30). Briefly, 50,000 AMs that were untreated, treated with poly (I:C) for four hours were spun down and washed once with cold PBS. The cells were resuspended in 50 μl cold ATAC-RSB-lysis buffer and incubated for 3 minutes. The ATAC-RSB-lysis buffer was immediately washed out with 1 mL ATAC-RSB buffer. The cell pellet was resuspended in 50 μl transposition mix and incubated for 30 minutes at 37 °C. The reaction was stopped by adding 2.5 μl pH 8 0.5 M EDTA. The Qiagen MiniElute PCR purification kit (Qiagen) was used to purify the transposed DNA. Purified DNA was amplified using the following condition: 72°C for 5 min, 98 °C for 30 s, and 13 cycles: 98 °C for 10s, 63 °C for 30 s, 72 °C for 1min. The amplified libraries were purified, size-selected, and the quality and quantity of libraries were assessed on 4150 TapeStation System (Agilent, CA). The pair-ended sequencing of DNA libraries was performed on an Illumina NovaSEQ6000 platform.

### ChIP-seq and Omni-ATAC-seq data analysis

The ChIP-seq and Omni-ATAC-seq data were analyzed as described before (29). Briefly, raw sequencing reads (average 40-80 million reads, 2 biological replicates for each treatment) were aligned to the hg38 reference genome using Bowtie2 with very-sensitive and -x 2000 parameters. The read alignments were filtered using SAMtools to remove mitochondrial genome and PCR duplicates. Peaks were identified by MACS2 with the q-value cut-off of 0.05 and the sequencing data was displayed using IGV.

### CRISPR plasmids constructions

The sgRNA guides targeting *TNF* enhancers were designed using CRISPICK (31,32) and compared selections of sgRNA guides with another designing tool CRISPOR (33). We analyzed and ranked potent off-target sites with Benchling. We selected sgRNA guides with high on-target scores and low off-target scores. Each of four sgRNA sequences targeting the same enhancer was cloned into LentiCRISPRv2GFP (Addgene, Plasmid # 82416), LentiCRISPRv2-mCherry (Addgene, Plasmid #99154), LentiCRISPRv2-BFP, or LentiCRISPRv2-THY1.1 vectors via the BsmBI cloning site. LentiCRISPRv2-THY1.1 vector was modified by replacing GFP gene in LentiCRISPRv2-GFP with the gene encoding THY1.1 using the SacII and BamHI restriction sites. Polymerases, restriction and modification enzymes were obtained from New England Biolabs (Beverly, MA).

### Construct the enhancer libraries using the lentiMPRA pLS-SceI vector

Enhancers were prepared and cloned into the lentiMPRA pLS-SceI vector as described (34) with modifications to allow the cloning of longer PCR-amplified enhancer fragments and sequencing barcode DNA and RNA using the Nova-seq multiplexing platform. PCR products of enhancers were amplified in a mix of PfuUltra DNA Polymerase (600385-51, Agilent Technologies, Santa Clara, CA) and *Taq* DNA polymerase (M7123, Promega, Madison, WI) at the ratio 1:10 using primers listed in Table S1. Read 1 and Read 2 sequencing primers (sequences listed in Table S2) were added PCR fragments by T/A ligation using NEBNext Ultra II Ligation Module (E7595, New England Biolabs, Ipswich, MA). Spacers and minimal promoter were added to sequencing primer-ligated enhancers using the primers 5BC-AG-f02 and 5BC-AG-r02 (Table S2). EGFP overhang and 15-bp random barcodes were added using 5BC-AG-f02 and 5BC-AG-r02 primers as described (24). Enhancer libraries were cloned into the lentiMPRA pLS-SceI vector (Addgene, Plasmid #137725) through Gibson assembly (E5510S, New England Biolabs, Ipswich, MA) according to manufacturer’s instructions.

For measuring enhancer activity using MPRA, 200-bp enhancer fragments were amplified using primers listed in Table S2. Read 1 and Read 2 sequencing primers, spacers, minimal promoter, EGFP overhang and 15-bp randomly selected barcodes were added to the enhancer fragments as above described. Enhancers with scan mutations were prepared as described (35). Briefly, linker-scanning mutations were introduced into the enhancer by PCR. Each linker-scanning mutation is 20 bp in length and lies within the context of the enhancer. PCR primers containing a 20 bp sequence (GTCAAGAGGTCGAGTCAAGA) that does not contribute to enhancer activity (25) were used to produce a series of PCR fragments that contain mutated sequence 20 bp at a time. The mutated sequences were joined with the rest of wild type sequences to generate 200 bp long enhancers using overlapping PCR as described (34). Read 1 and Read 2 sequencing primers, spacers, minimal promoter, EGFP overhang and 15-bp random barcodes were added to the mutated enhancers as above described.

### Enhancer sequencing and barcode association

The mutant enhancers plasmid library was amplified with the following primers: P7_association_N7## and P5_association_N5## (Table S2). The library was amplified using the following condition: 98 □ for 1 min, then 15 cycles: 98 □ for 15 s, 60 □ for 20 s, 72 □ for 3 min. And 72 □ for 5 min. The DNA product was purified from the gel slice using the PCR cleanup and gel extraction kits (740609.50, Takara Bio, Kusatsu, Shiga). The purified DNA was cleaned using AMPure XP (A63881, Beckman, Brea, CA) according to the manufacturer’s instructions. The quality and quantity of libraries were assessed on the 4150 TapeStation System (Agilent, CA). The pair-ended sequencing of DNA libraries was performed on an Illumina MiSeq platform by using MiSeq Reagent Kit v2 300-cycles PE (MS-102-2002, San Diego, CA): 138 cycles for read1, 15 cycles for custom index1, 8 cycles for index2 and 138 cycles for read2.

### Barcode sequencing and counting

The cells were washed with PBS three times, and genomic DNA and total RNA were extracted using a DNA/RNA mini kit (Qiagen) according to the manufacturer’s instructions. LentiMPRA barcoded RNA-seq and DNA-seq libraries were constructed according to the published method with P7 and P5 counting primer modifications (24). Briefly, the original custom read 1 and read 2 primers were replaced by the read 1 and read 2 primers listed in the Table S2. An i7 index was added to the 5’ of read 2. DNA with inserted barcodes or reverse transcription products containing barcodes were amplified with the modified P7_counting and P5_counting primers listed in the Table S2. The preliminary PCR reaction was set up for each sample to determine the number of cycles at which the amplification nearly plateaus for each sample as described (24). P5 and P7 primers listed in the Table S2 were used to amplify PCR products for sequencing. The final PCR product was purified using AMPure XP (A63881, Beckman, Brea, CA) according to the manufacturer’s instructions. The quality and quantity of libraries were assessed on 4150 TapeStation System (Agilent, CA). The pair-ended sequencing of DNA libraries was performed on an Illumina NovaSEQ6000 platform; standard sequencing 151+8+10+151, which is read1+index1+index2+read2 cycles. Barcodes were extracted from read 1 and read2, and UMI were extracted from read1 raw sequencing data using umi-tools (version: 1.1.1). For longer enhancers (>200 bp), the number of DNA and RNA containing manually selected barcodes were sequenced using Illumina NovaSEQ6000 or measured by qPCR using primers listed in Table S1.

### MPRA processing pipeline

Barcodes were associated with enhancer sequences, and the number of bar codes in the RNA and DNA samples was counted using the software MPRAflow as described in the published bioinformatics workflows (24). We analyzed the NGS sequencing data on Linux. The codes were downloaded from https://docs.conda.io/en/latest/miniconda.html, and the MPRAflow was downloaded from https://github.com/shendurelab/MPRAflow.git. For the barcode association, the code is “nextflow run association.nf --fastq-insert “R1_001.fastq.gz” --fastq-insertPE “R3_001.fastq.gz” --design “ordered_candidate_sequences.fa” --fastq-bc “R2_001.fastq.gz”“. For the barcode counting, the code is “nextflow run count.nf --dir “bulk_FASTQ_directory” --e “experiment.csv” –design “ordered_candidate_sequences.fa” –association “dictionary_of_candidate_sequences_to_barcodes.p”“.

### Lentivirus production and transduction

The 10cm dishes were coated with 4 mL l0 μg/mL poly D lysine (Sigma, P0899) for 5 minutes at room temperature in H_2_O. Plate cells at 2-3 × 10^6^ HEK293T cells/dish in DMEM (10% FBS, but no antibiotics). Twenty-four hours later, HEK293T cells were transfected with 10 μg of pLS-SceI-BE plasmid or four color bicistronic sgRNA guides LentiCRISPRv2 plasmids, 9 μg PΔ8.9 and 1 μg VSV-G using CaCl_2_. Seventy-two hours after transfection, the supernatants were collected and filtered with a 0.45 μm filter using procedures described (36).

THP-1 cells were cultured in RPMI 1640 medium plus 10% FBS, 100 units/mL penicillin, 100 μg/mL streptomycin and 2mM beta-mercaptoethanol in a humidified 37°C, 5% CO_2_ incubator. The 1×10^6^ THP-1 cells were seeded into one well of a 6-well plate with 10 mL lentivirus medium. The polybrene was added to each plate at the final concentration of 8 μg/mL and the HEPES was added to each plate at final concentration of 25 mM. Lentivirus supernatants were added to each plate. The plates were wrapped with parafilm and centrifuged at 2,500 rpm for 90 minutes at room temperature. The supernatant was removed by aspiration and 2 mL fresh medium per well was added. The spin infection step was repeated at the next day and the day after for a total of three spin infections. Two days after the last spin infection, the cells for four color sgRNA guides deletion that expressed BFP, GFP, RFP, and Thy1.1 were FACS-sorted. For the reporter assay, the transduced THP-1 cells can be used for further analysis.

### PMA maturation and poly (I:C) stimulation

THP-1 cells express low levels of pro-inflammatory cytokines upon poly (I:C) stimulation. These cells can be matured to become robust pro-inflammatory cytokine-producing cells by Phorbol 12-Myristate 13-Acetate (PMA, Sigma-Aldrich). Transduced-THP-1 cells were incubated with PMA (200 ng/mL) for three days. The matured THP-1 cells were not treated or treated with 20 μg/mL poly (I:C) (4287, R&D Systems, Minneapolis, MN) for four additional hours before the cells were collected for analysis.

### Transcription factor binding motif analysis

The transcription factor binding motif analysis were conducted by using R package TFBSTools (version 1.36.0) (37) and Biostrings (version 2.66.0) (38). TF binding motif matrices from the JASPAR 2022 database (39) were used in the analysis.

### qPCR analysis

The untreated or poly (I:C)-stimulated THP-1 cells were collected; genomic DNA and total RNA were extracted using a DNA/RNA mini kit (Qiagen) according to the manufacturer’s instructions. Quantitative PCR was performed in a QuantStudio 7 Flex Real-Time PCR System (ThermoFisher, MA). The sequences of qPCR primers are listed in Table S1. Relative mRNA amounts were calculated as follows: Relative mRNA or DNA amount = 2^[Ct(Sample)-Ct(*HPRT*)]^. The barcode reporter activity was measured as the ratio of RNA and DNA.

## Data availability

All raw and processed sequencing data generated in this study have been submitted to the NCBI Gene Expression Omnibus (GEO; https://www.ncbi.nlm.nih.gov/geo/) under accession number GSE226158.

## Statistical analysis

The nonparametric Mann-Whitney U test or two-tailed student’s *t*-test was used to determine significant differences between the two treatment groups.

## Acknowledgement

We thank laboratory members for thoughtful discussions. We are grateful to Marlene Gallegos Sanchez for technical assistance. We are thankful to Dr. Robert J. Mason for collection of the human alveolar macrophages. We are grateful to Dr. Bifeng Gao and the staff of the Genomics Shared Resource Facility at the University of Colorado Cancer Center for Next-Generation Sequencing.

## Author contributions

Conceptualization, H.H. and J.G.; methodology, J.G., Y.L., H.W.C., J.S.B. and H.H.; investigation, J.G., Y.L., D.Z., X.G. and K.M.; visualization, J.G. and Y.L.; funding acquisition, H.H.; supervision, H.H.; writing-original draft, J.G. and H.H.; writing-review & editing, J.G., H.W.C., J.S.B. and H.H.

