## Supplementary material for "Enhancers that regulate *TNF* gene transcription in human macrophages in response to TLR3 stimulation": Fig. S1: Figure S1.pdf

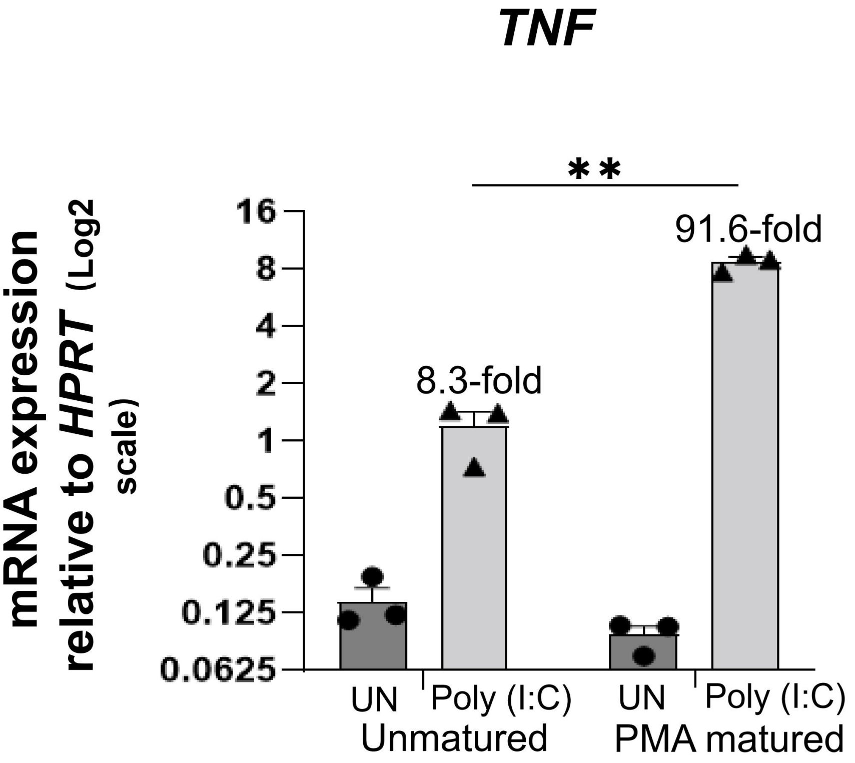


**Figure S1.** PMA matured THP-1 transcribed high levels of *TNF* mRNA in response to poly (I:C) stimulation. THP-1 cells were incubated without or with PMA for three days. The unmatured or PMA matured THP-1 cells were not stimulated (UN) or stimulated with 20 μg/mL Poly (I:C) for four additional hours before the cells were collected for qPCR analysis. *P* values were calculated using two-tailed student's *t* test. Data represents mean ± SEM of three samples. ** $P < 0.01$.
