## Supplementary material for "Enhancers that regulate *TNF* gene transcription in human macrophages in response to TLR3 stimulation": Fig. S2: Figure S2.pdf

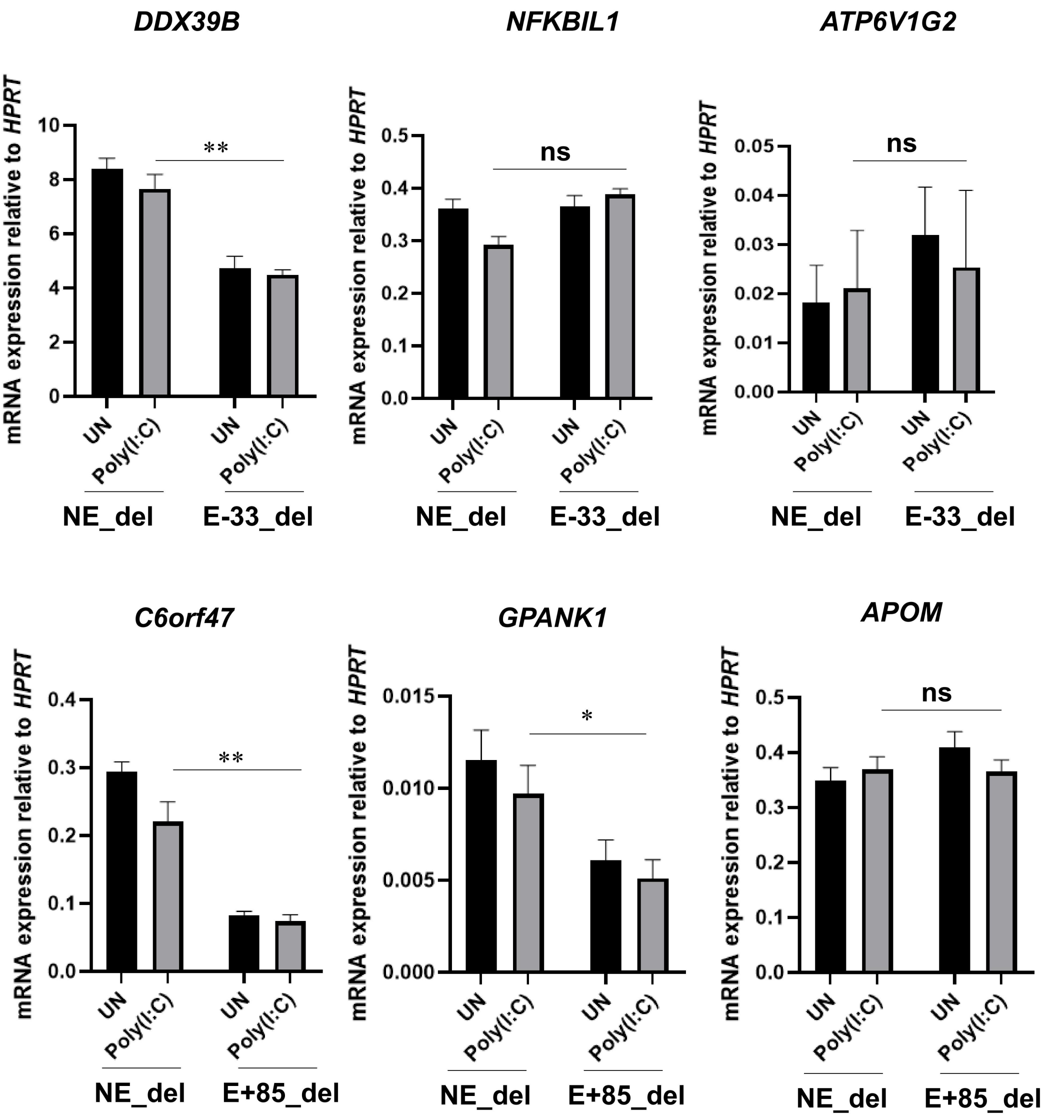


**Figure S2. The E-33 is required for *DDX39B* gene transcription and the E+85 is required for *C6orf47, GPANK1* gene transcription. A.** THP-1 cells were transduced with E-33 GFP, BFP, RFP, and Thy1.1 sgRNA guides (4c sgRNA guides). *DDX39B*, *NFKBIL1* and *ATP6V1G2* mRNA expression in the FACS-sorted sgRNA guides transduced cells was measured by qPCR. **B.** THP-1 cells were transduced with E+85 GFP, BFP, RFP, and Thy1.1 sgRNA guides. *C6orf47*, *GPANK1* and *APOM* mRNA expression in the FACS-sorted sgRNA guides transduced cells was measured by qPCR. *P* values were calculated using two-tailed student's *t* test. Data represent mean ± SEM of four transduced samples. * $P < 0.05$; ** $P < 0.01$.
